# Real-time monitoring of pediatric high-grade glioma invasion using organotypic brain cultures reveals a developmentally dependent sensitivity

**DOI:** 10.64898/2025.12.24.696338

**Authors:** Yoav Biala, Ramakrishna Sai kumar Aravapalli, Hala Shalata, Zakhariya Manevitch, Oded Behar

## Abstract

High-grade gliomas (HGGs) constitute a considerable portion of pediatric brain tumors, typically arising in the supratentorial cortex (median age, 13 years) or infratentorial/midline regions (median age, 7 years). About half are diffuse midline gliomas (DMGs), whose aggressive invasiveness contributes significantly to their lethality. Monitoring tumor behavior in its native environment, especially invasion, remains challenging. Although mouse models are used in glioma research, pediatric HGGs present unique difficulties. Multiphoton microscopy allows cortical invasion studies, but DMG analysis is hindered by deep anatomical locations and rapid mouse aging, which limits relevance to developmental stages.

To overcome these barriers, we established a novel platform combining direct implantation of pontine DMG and cortical HGG cells into organotypic *ex vivo* brain slice cultures with spinning-disc confocal time-lapse imaging. This method enables accelerated, real-time monitoring of tumor invasion across multiple slice cultures under conditions that recapitulate some of the key features of the native microenvironment. Using this system, we demonstrated that the developmental stage of brain tissue strongly influences pontine DMG invasion and validated the platform’s utility for evaluating anti-invasive therapies.

## Introduction

High-grade gliomas (HGGs) constitute a considerable portion of pediatric brain tumors and are typically fatal within two years of diagnosis (Cosnarovici *et al*, 2021). These tumors develop in two main regions: the supratentorial cerebral cortex and the infratentorial regions, including the cerebellum, brainstem, midbrain, thalamus, and spine (Mackay *et al*, 2017). Nearly half of the pediatric HGGs (pHGGs) arise in midline locations, particularly the thalamus and pons. They are classified as diffuse midline gliomas (DMGs) with pontine DMG, also known as diffuse intrinsic pontine glioma (DIPG). The age of onset varies by location: pontine and thalamic DMGs have a median age of diagnosis around age 7, while cortical gliomas have a median age of diagnosis at age 13. In contrast, adult HGGs predominantly occur in the frontotemporal lobes (Mackay *et al*, 2017; Juratli *et al*, 2018).

Supratentorial and infratentorial pHGGs also differ in their molecular characteristics. A notable example is the lysine-to-methionine mutation at residue 27 in histones H3.1 (H3.1K27M) and H3.3 (H3.3K27M), found in approximately 85% of pontine DMG cases in young children. In contrast, H3.3G34R/V mutations typically occur in cerebral hemisphere tumors of adolescents and young adults (Juratli *et al*, 2018; Mackay *et al*, 2017). The correlation between tumor anatomical position, age preference, and specific mutations suggests a unique tumor-microenvironment compatibility. This correlation further indicates that studying these pediatric tumor types should preferably be conducted in their appropriate anatomical and developmental contexts to properly understand the tumor-microenvironment cross-talk. These specific conditions may be challenging to replicate using current experimental systems.

The fatal nature of all HGGs, including pediatric cases, stems at least in part from their diffused invasion throughout the brain parenchyma, which resists therapeutic intervention. While various methods exist to study adult glioma invasion using slice cultures and *in vivo* modeling with live imaging, these approaches have limitations. Although *in vivo* systems provide the most physiologically relevant context, they present challenges for monitoring tumor invasion, particularly when considering personalized medicine applications. DMG cases pose additional challenges due to their anatomical location, which prevents multiphoton microscopy imaging *in vivo*. Even in slice culture studies, continuous high-resolution monitoring of tumor behavior throughout experiments has proven difficult (Haydo *et al*, 2023; Mann *et al*, 2023; Pencheva *et al*, 2017).

Studying pediatric gliomas in mouse models presents unique challenges due to mice’s accelerated development compared to humans. When using allograft or patient-derived xenograft (PDX) systems, tumor cell injection before one month of age is challenging. Thus, most studies inject tumor cells at 4-6 weeks of age (Tsoli *et al*, 2019; Barron *et al*, 2025). Direct conversion between human and mouse developmental stages is challenging(Pathania *et al*, 2017)(Pathania *et al*, 2017). However, given the rapid maturation of mice, pediatric tumor progression in these models predominantly occurs within adult-like microenvironments. This developmental mismatch poses a significant limitation for studies investigating the role of the pontine microenvironment in DIPG formation and progression during early development, underscoring the critical need for complementary experimental approaches.

In this study, we developed an organotypic brain slice culture system that enables high-resolution monitoring of pediatric tumor invasion over time in both pontine and cortical tissue. We demonstrated that the system maintains tissue viability and structural integrity, that DIPG invasion is sensitive to host tissue age, and that the platform is suitable for drug screening in a physiologically relevant environment.

## Results

### Characterization of slice culture and tumor cells in the pons and cortex

Organotypic slice cultures that incorporate tumor cells typically use one of two methods: either plating tumors on top of the slices (Haydo *et al*, 2023; Chadwick *et al*, 2015; Sun *et al*, 2023; Decotret *et al*, 2023; Unable to find information for 16371064) (Pencheva *et al*, 2017) or implanting the tumor cells directly into the slices (Petterson *et al*, 2016; van Asperen *et al*, 2022). While implanting tumor cells into the slices is more technically challenging, especially for continuous monitoring purposes, this approach provides an environment more similar to the cells’ native conditions. Therefore, we chose to implant tumor cells directly inside tissue (Figure 1a). To monitor the behavior of DIPGs in their native microenvironment, small tumor spheroids were microinjected into the center at the Z-axis of relevant brain slices. Although the morphology of initial tumor formation in patients remains unknown, we selected spheroid implantation for its practical advantages: enhanced initial cell survival and simplified tracking of invasive cells migrating from the spheroid mass. To enable visualization of tumor cells, we labeled them with lenti-GFP. To minimize tissue damage during insertion, we used glass pipettes with a 75-100 μm diameter. To test the characteristics of the invasion of DIPG (SU-DIPGXIII) in the pons or cortical HGG (SU-pcGBM2) tumor cells in the cortex, we injected the tumor spheres a day after slice preparation and maintained them for up to 7 days. After fixation, the slices were sectioned and evaluated using H&E staining (Figure 1b). Overall, the integrity of the tissue was maintained. To test the state of the tissue, we also used antibodies to visualize the tumor cells as well as neurons, astrocytes, and microglia in the pons (Figure 1c) or the cortex (Figure 1d).

**Figure 1:**
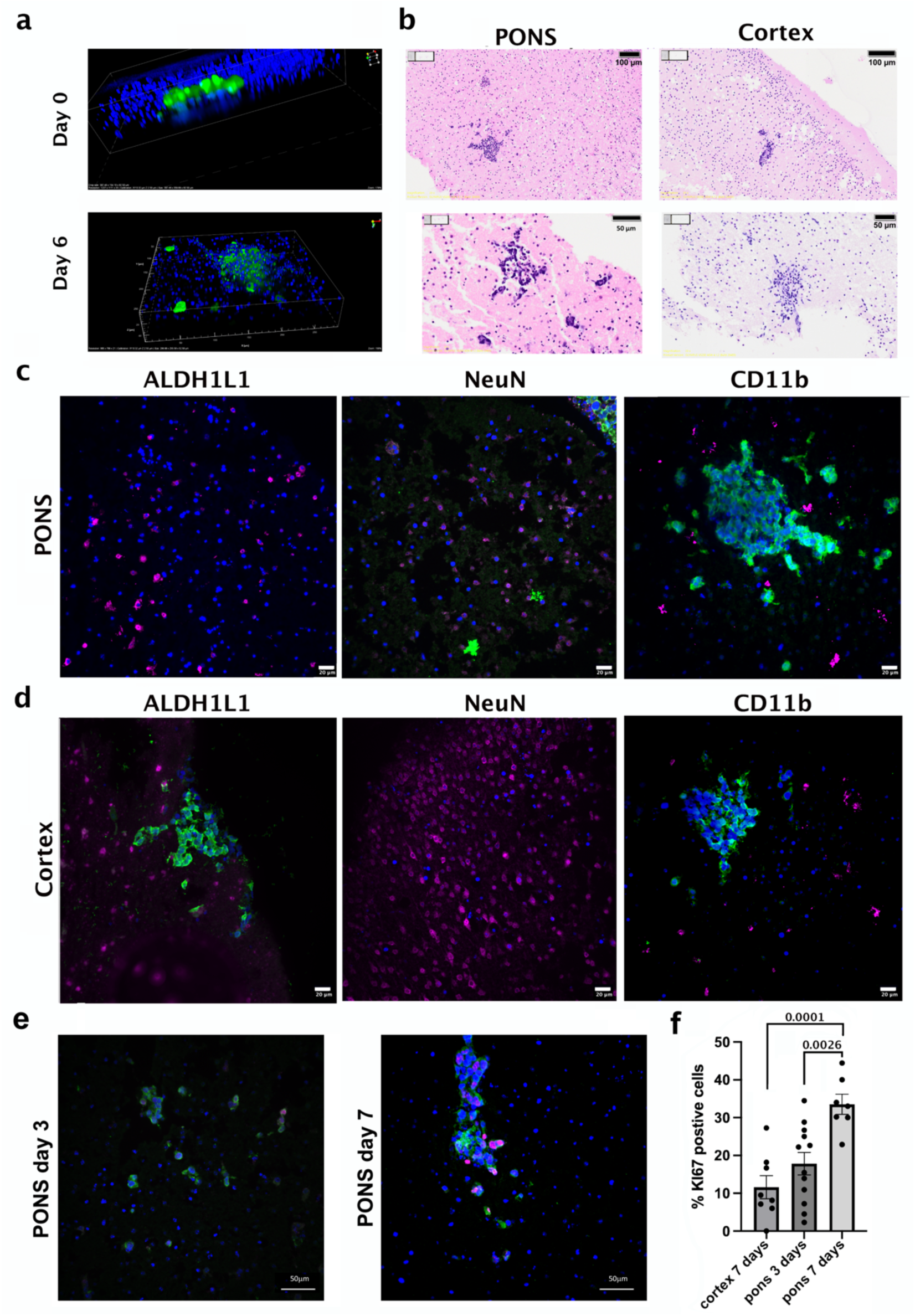
Characterization of pontine and cortical gliomas in organotypic slice cultures. a) GFP-expressing SU-DIPGXIII tumor spheres (green) embedded within brain slices at 2 hours post-injection (upper panel) and after 7 days in culture (lower panel). Nuclei are labeled with Hoechst (blue). b) H&E staining of paraffin sections from one-week-old pontine slices injected with SU-DIPGXIII (left images) and cortical slices injected with SU-pcGBM2 (right images). Scale bars: 100 μm (upper images), 50 μm (lower images). c) Immunostaining of paraffin sections from one-week-old pontine slices showing nuclei in blue, tumor cells in green, and different cell types in magenta (ALDH1L1 for astrocytes, NeuN for neurons, and CD11b for microglia/macrophages). Scale bar: 20 μm. d) Immunostaining of paraffin sections from one-week-old cortical slices showing nuclei in blue, tumor cells in green, and different cell types in magenta (ALDH1L1 for astrocytes, NeuN for neurons, and CD11b for microglia/macrophages). Scale bar: 20 μm. e, f) SU-DIPGXIII cells were injected into pontine and cortical slices obtained from one-week-old mice. Paraffin sections were analyzed at days 3 and 7 post-injection. e) Representative images of Ki67 staining in tumor cells at days 3 and 7 in the pons. Scale bar: 50 μm. f) Quantification of Ki67-positive tumor cells showing increased proliferation over time in the pons, with higher proliferation rates compared to the cortex. (n=7-12 mice; Mann-Whitney two-tailed test).

**Figure 2:**
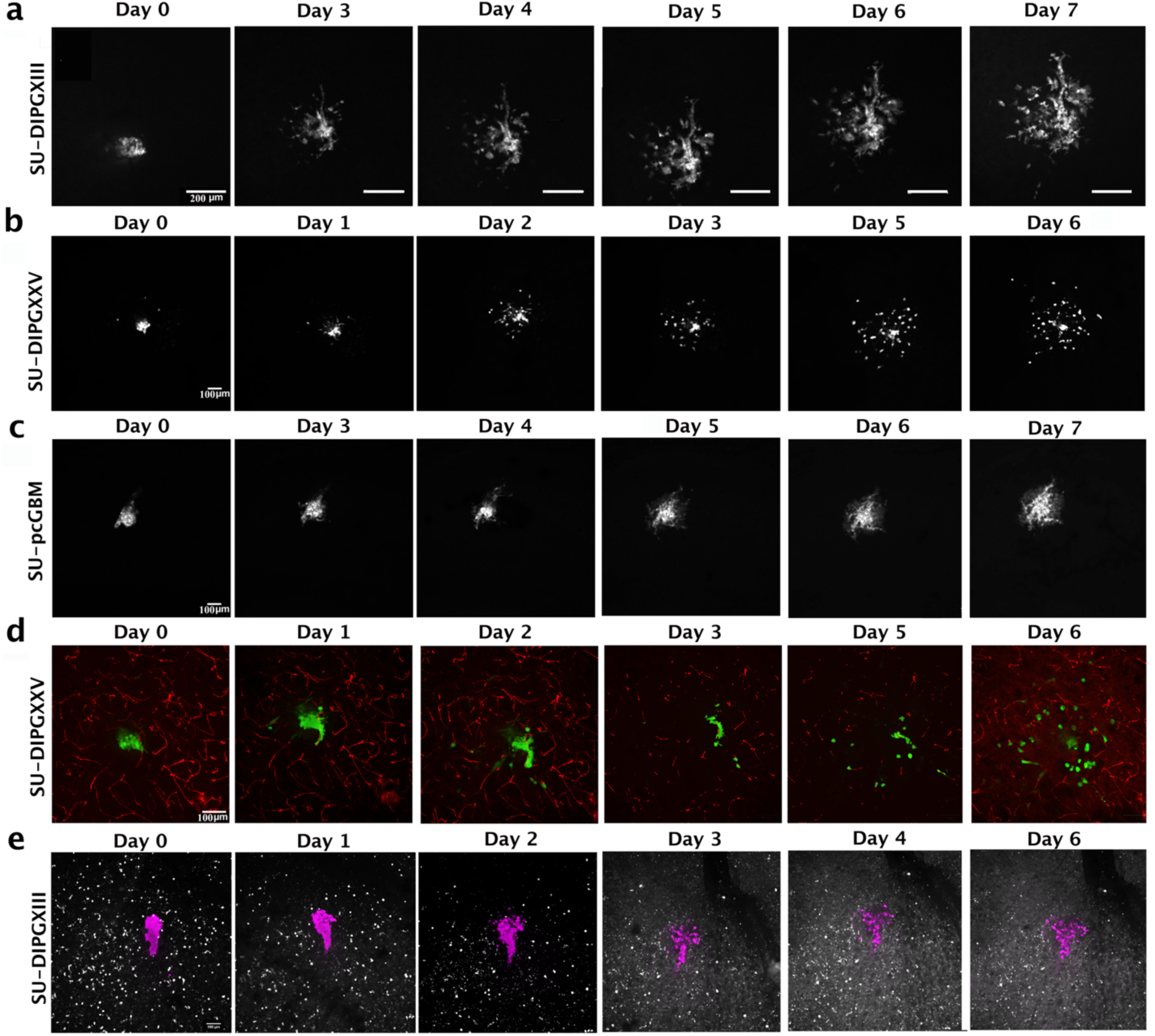
Longitudinal monitoring of tumor behavior in tissue a) Time-lapse imaging of a representative SU-DIPGXIII tumor sphere in the pons at 2 hours post-injection and on subsequent days. Scale bar: 200 μm. b) Time-lapse imaging of a representative SU-DIPGXXV tumor sphere in the pons at 2 hours post-injection and on subsequent days. Scale bar: 100 μm. c) Time-lapse imaging of a representative SU-pcGBM2 tumor sphere in the cortex at 2 hours post-injection and on subsequent days. Scale bar: 100 μm. d) Time-lapse imaging of a representative SU-DIPGXXV tumor sphere in pontine slices with pre-labeled blood capillaries (IB4-Cy3) at 2 hours post-injection and on subsequent days. Scale bar: 100 μm. e) Time-lapse imaging of a representative SU-DIPGXIII tumor sphere in pontine slices from Cx3Cr1-GFP mice (labeled microglia/macrophages) at 2 hours post-injection and on subsequent days. Scale bar: 100 μm.

Tumor cells persist, and their numbers appear to increase over time. Additionally, cells detach from the spheres and invade the surrounding tissue. To better assess the proliferative activity of tumor cells, we performed Ki67 staining on tissue sections at days 3 and 7 post-injection (Figure 1e, f). DIPG tumor cells progressively proliferate and migrate within the tissue, demonstrating their adaptive capacity. Notably, DIPG cell proliferation increases with extended duration in the slice cultures (Figure 1e). Furthermore, DIPGs implanted in the cortex exhibited reduced proliferation at day 7 compared to the same tumors implanted in the pons, underscoring the critical role of the pontine microenvironment in supporting DIPG growth. These findings are consistent with our previous observations (Zats *et al*, 2022).

### 3D time-lapse imaging of organotypic slice cultures

The invasive growth pattern of pediatric and adult high-grade gliomas is a primary factor contributing to their lethality. To develop effective therapeutic interventions, it is essential to monitor this invasive behavior under conditions that closely mimic the patient environment, as such systems provide opportunities to test potential inhibitors of tumor invasion. This requires the ability to monitor tumor cell invasion at high resolution over extended time periods.

Longitudinal analysis of tumor cell invasion in slice cultures necessitates the capability to repeatedly return to identical locations in 3D space and image the same tumor cells at multiple time points. To achieve high-resolution monitoring of multiple tissue slices, we adapted confocal spinning disc microscopy. Imaging tumor cells in organotypic slice cultures is challenging due to the working distance required by the microscope objective. To minimize this working distance, we developed a specialized chamber using 3D printing in which tissue slices are positioned on a transparent membrane in close proximity to a glass-bottom plate (Supplementary Figure 1). To characterize the invasive properties of DIPG cells within the pons and cortical tumor cells within the cortex, we performed multi-day imaging of identical tumor cell populations to track their invasion into surrounding tissue (Figure 3a-c).

**Figure 3:**
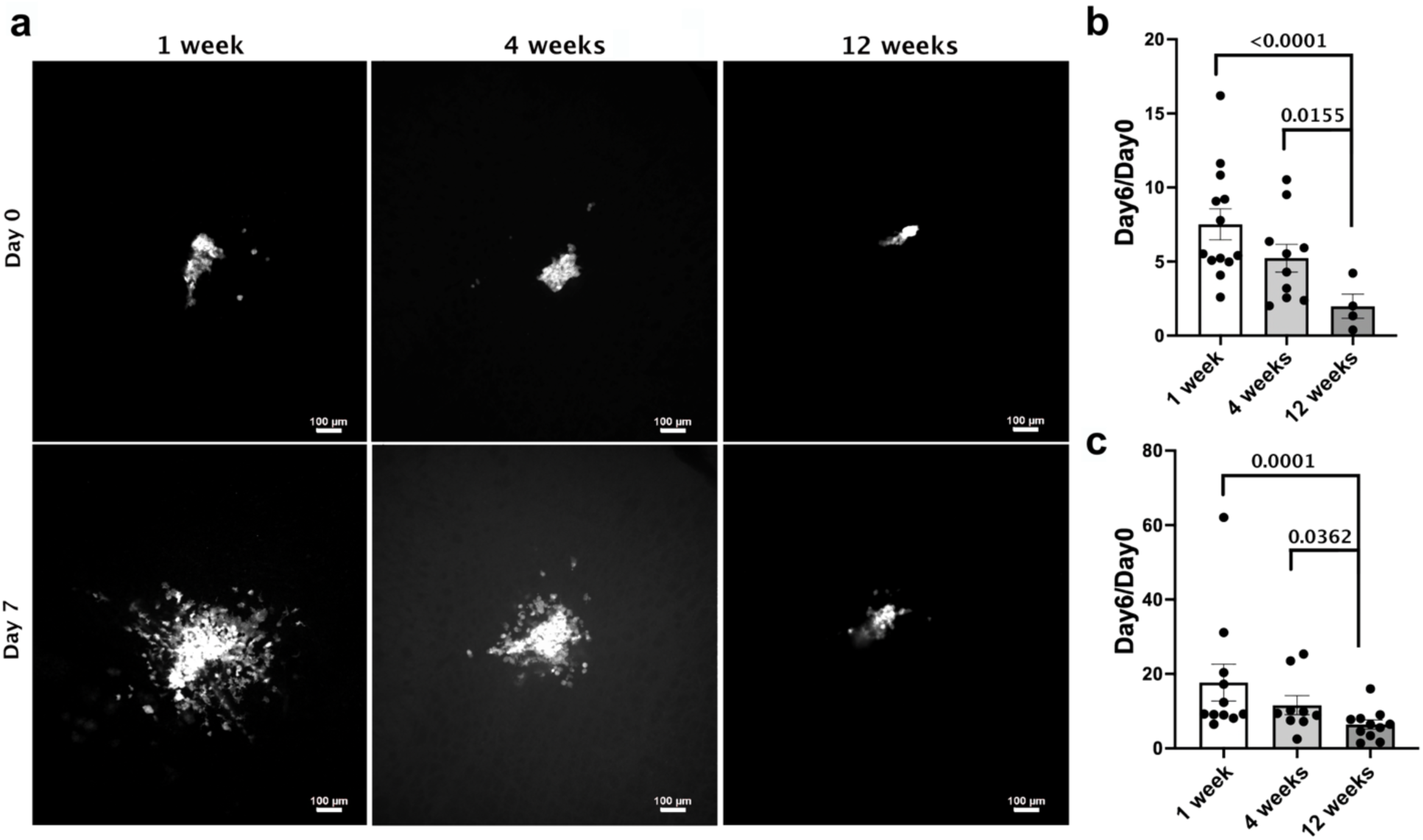
DIPG tumor invasion decreases with increasing mouse age a) Representative images of SU-DIPGXIII tumors at the time of injection and 7 days post-injection in pontine slices prepared from 1-, 4-, and 12-week-old mice. Scale bar: 100 μm. b) Quantification of SU-DIPGXIII tumor invasion in slices prepared from 1-, 4-, and 12-week-old mice. (n=5-13 mice; Mann-Whitney two-tailed test). c) Quantification of SU-DIPGXXV tumor invasion in slices prepared from 1-, 4-, and 12-week-old mice. (n=9-11 mice; Mann-Whitney two-tailed test).

To examine interactions between DIPG cells and resident tissue cells, we performed daily imaging using two complementary approaches: (1) GFP-labeled SU-DIPGXXV tumor cells in brain slices pre-labeled with Isolectin-B4 (IB4) conjugated to Cy3, a standard marker for visualizing blood capillaries (Figure 3d, Supplementary Figure 2); and (2) RFP-labeled SU-DIPGXIII tumor cells in brain slices from Cx3Cr1-GFP mice to visualize microglia/macrophages (Figure 3e). Both experimental strategies demonstrate our methodology’s capacity to simultaneously monitor tumor cells and key microenvironmental components. Notably, SU-DIPGXXV tumor cells appear to migrate along blood capillaries preferentially, a behavior consistent with high-grade gliomas (HGGs) in adults(Krusche *et al*, 2016; Mair *et al*, 2018)(Krusche *et al*, 2016; Mair *et al*, 2018).

### The impact of mouse age on DIPG tumor invasion potential

Most experiments designed to model pediatric tumor behavior utilize mice aged 5-8 weeks. To assess how the developmental stage of the brain microenvironment influences tumor invasion, we implanted SU-DIPGXIII and SU-DIPGXXV tumor spheres into brain slices harvested from mice at 1, 4, and 12 weeks of age and compared their invasive behavior. Sections were imaged shortly after implantation into the pons and again 6 days later (Figure 4). The results demonstrate that DIPGs, which were derived from young patients, exhibit more aggressive invasion in younger pontine tissue, highlighting the advantage of our system for evaluating age-specific properties of pediatric tumors.

**Figure 4:**
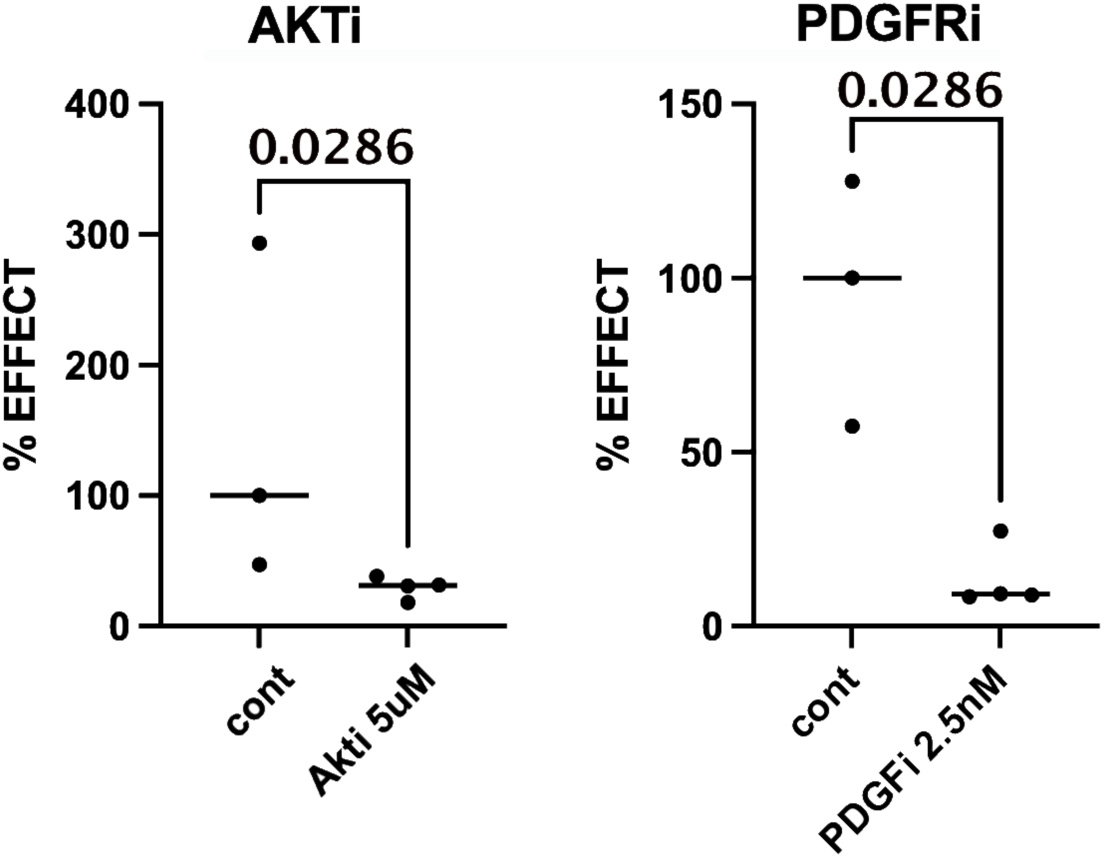
Impact of targeted inhibitors on invasive growth in the pons Quantification of SU-DIPGXIII tumor invasion in one-week-old pontine slices treated with or without AKT and PDGFR inhibitors. (n=3-4 mice; Mann-Whitney one-tailed test).

The ability of a given drug to effectively target a specific tumor in a patient is central to personalized treatment strategies. The most straightforward approach to assessing drug efficacy involves testing patient-derived tumor cells in vitro. However, tumor cells within their native microenvironment can behave quite differently from those cultured in isolation. To determine whether our method can be used to evaluate drug sensitivity, we examined the effects of two inhibitors—JNJ-10198409, a PDGFR inhibitor, and MK-2206, an AKT inhibitor—on the invasive growth of DIPG cells using the SU-DIPGXIII model in organotypic slice cultures. Both inhibitors nearly completely suppressed the invasive capacity of SU-DIPGXIII cells in slice culture (Figure 5), thus demonstrating the utility of this method for testing tumor cell drug sensitivity.

## Discussion

The ability to develop new treatments and test their suitability for individual patients depends on the model system’s capacity to provide fast and reliable information for treatment evaluation. In this study, we demonstrate a slice culture system that enables high-resolution monitoring of HGGs in relevant brain positions and developmental stages.

### Tumor monitoring using a slice culture system

Pediatric HGGs, including DMGs, are highly invasive tumors with poor prognoses despite more than 250 clinical trials over three decades. The urgent need for effective treatments has been hampered by inadequate preclinical models that fail to accurately represent the dynamic tumor microenvironment (TME) and tumor behavior in native brain environments.

Traditional approaches rely on 2D and 3D cultures, which lack the complexity of the tumor microenvironment. While 3D cultures and co-culture systems have improved physiological relevance, they still cannot fully recapitulate the dynamic tumor-brain interface. Mouse in vivo models offer distinct advantages, particularly in providing microenvironmental complexity that closely resembles physiological tumor conditions. However, these systems suffer from two critical limitations. First, the accelerated pace of mouse brain development relative to humans means that tumor behavior is studied in a more mature microenvironment than is developmentally appropriate for modeling pediatric tumors. Second, limited imaging accessibility, especially for deep midline structures, prevents detailed characterization of invasive behavior, a hallmark of HGGs.

Although combining in vivo modeling with ex vivo imaging addresses some imaging constraints, the accelerated developmental timeline of the mouse system remains a fundamental limitation.

Slice culture models represent a compromise between *in vitro* and *in vivo* approaches, offering an opportunity to test HGG responses in a microenvironment most similar to *in vivo* conditions. These systems can theoretically enable monitoring of tumor behavior within its native microenvironment, though they are temporally limited. However, technical challenges have historically limited the ability to monitor tumor behavior over time at high resolution. The poor concordance between preclinical studies and clinical trials for H3K27M-altered DMG/DIPG highlights these modeling limitations and underscores the need for improved experimental systems.

Here, we present a novel *ex vivo* approach combining direct implantation of DIPG and cortical HGG cells into organotypic brain cultures with spinning disc confocal time-lapse imaging. This platform enables high-resolution, real-time visualization of tumor invasion over multiple days within a physiologically relevant microenvironment, allowing simultaneous monitoring of tumor cells, blood vessels, microglia, and other cell types.

Our findings reveal key insights: DIPG cells seem to largely follow blood vessels during invasion, and tumor behavior is highly sensitive to developmental stage, with significantly reduced invasive capacity in tissue from older mice. Despite originating from human patients and mouse host tissue, tumor cells recognize appropriate brain regions and show enhanced proliferation and invasion when placed in age-matched, anatomically correct locations. These results highlight the inherent limitations of studying pediatric tumors in standard mouse models with accelerated aging.

Compared to existing models, our *ex vivo* system preserves native tissue architecture while enabling real-time, high-resolution imaging without confounding developmental effects. This addresses critical gaps by providing a physiologically relevant platform for mechanistic studies and therapeutic evaluation that bridges *in vitro* and *in vivo* approaches.

The platform’s versatility across tumor types and brain regions, combined with potential expansions incorporating patient-derived samples, immune components, and pharmacological screening, positions it as a valuable tool for personalized therapeutic testing. By enabling real-time monitoring of tumor behavior in developmentally appropriate tissue, this approach enhances mechanistic understanding and offers new opportunities for preclinical testing of anti-invasive therapies, potentially accelerating translation to improve outcomes for children with these devastating tumors.

## Methods

Animals: C57BL/6 mice were obtained from ENVIGO (Rehovot, Israel). Animal handling adhered strictly to national and institutional guidelines for animal research and was approved by the Ethics Committee of the Hebrew University.

### Patient-derived tumor cell cultures

All cultures were maintained as neurospheres in tumor stem medium (TSM) consisting of 50% Neurobasal(-A)/50% DMEM/ F12, supplemented with B27TM(-VitA) (all from Thermo Scientific, Waltham, MA, USA) and growth factors as described previously^16^ and in Supplementary methods.

## Organotypic slice culture preparation

<u>Dissection and slice preparation</u> are detailed in the supplementary methods.

### Injection of cancer cells

One day after initial incubation of the slices, tumor spheres were injected using glass capillaries pulled with a Sutter P-97 puller (Sutter Instrument, Novato, CA, USA) equipped with a 3 × 3 mm box filament using the following parameters: pressure 500, heat 614, pull 0, velocity 150, time 0. Capillaries were cut at an angle with a scalpel to create a beveled tip opening, facilitating tissue penetration. Spheres of SU-DIPGXIII, SU-DIPGXXV, or SU-pcGBM2 were injected at multiple discrete locations within each slice, targeting the mid-depth (Z-position) of the tissue. Spheres were imaged at 2 hours post-injection and subsequently either daily or only at the experimental endpoint (day 6 or 7). Cells were imaged using a Nikon ECLIPSE Ti2 confocal inverted microscope with an incubator to acquire high-resolution fluorescence images (Nikon Corporation, Tokyo, Japan). The microscope is equipped with an Orca FusionBT monochrome camera (Hamamatsu Photonics, Hamamatsu, Japan) and a CSU-W1 spinning disk unit with a 50 µm pinhole (Yokogawa Electric Corporation, Tokyo, Japan). The system was integrated with 405, 488, 561, and 638 nm LH+ lasers and a 10x CFI Plan Apochromat lambda 0.45 N.A., 20x CFI Plan APO VC 0.75 N.A., and 40x Plan Apochromat 0.95 N.A. Images were analyzed using either NIS or FIJI software.

### Slice preparation and Immunofluorescence

Details regarding the Slice preparation, immunofluorescence, and antibody use are in the supplementary methods.

## Use of an LLM

A large language model (Claude, Anthropic) was used to improve the English and grammar of the manuscript text.

## Statistical analysis

Values presented are mean ± SEM. P-value ≤ 0.05 was considered significant. Statistical analysis was performed using the non-parametric two-tailed or one-tailed Mann–Whitney U test.

## Acknowledgments

This work was supported in part by a grant from the The Cure Starts Now Foundation, Brooke Healey Foundation, Melina Michelle Edenfield Foundation, The Cure Starts Now Australia, The Cure Starts Now Canada, Reflections Of Grace Foundation, Yuvaan Tiwari Foundation, Cure Brain Cancer Foundation, Aubreigh’s Army Foundation 328, Aidan’s Avengers, Run DIPG, Musella Foundation, Love4Lucas Foundation, Whitley’s Wishes, Anna’s Bake Sale Foundation, The Ayla Foundation, The Isabella and Marcus Foundation, Love, Chloe Foundation, Lauren’s Fight for Cure, Robert Connor Dawes Foundation, Ryan’s Hope, The Gold Hope Project, Abby’s Corner Foundation, The DIPG/DMG Collaborative and Snapgrant.com.

We thank Vitali Belzer and Dr. Liat Peretz Zats for contributions to initial method development, Michelle Monje (Stanford University School of Medicine) for providing the cell models SU-pcGBM2, SU-DIPGXXV, and SU-DIPGX, and the Hebrew University 3D & Functional Printing Center for technical advice.

## Author Contributions

Y.B. established the method design, contributed to Figures 1-3, and helped prepare the manuscript. R.A. performed experiments in Figures 2-4 with significant help from H.S. Z.M. adapted the spinning disc microscopy system. O.B. designed all experiments, wrote the manuscript, and supervised the work.

## Competing Interests

There are no competing financial interests in relation to the work described.

## The Paper Explained

### PROBLEM

High-grade gliomas are aggressive brain tumors that account for 8-10% of pediatric brain cancers. Roughly half of these tumors develop in critical midline brain structures, including the pons (a region essential for vital functions like breathing and heart rate). These diffuse midline gliomas are particularly deadly because they invade surrounding brain tissue aggressively, making them impossible to remove surgically or treat effectively. A major obstacle in developing better treatments is the difficulty of studying how these tumors invade brain tissue in real time. Current mouse models face significant limitations: tumors located deep in the brain are difficult to visualize, and mice age rapidly, making it hard to study tumors in brain tissue that matches the developmental stage of pediatric patients. Without effective tools to observe and understand tumor invasion as it happens, researchers struggle to identify and test therapies that could stop these cancers from spreading.

### RESULTS

We developed a new experimental platform that allows researchers to watch pediatric glioma cells invade brain tissue in real time. Our approach combines two key techniques: first, we implant tumor cells directly into preserved brain slices taken from the pons (for midline gliomas) or cortex (for other high-grade gliomas); second, we use advanced time-lapse microscopy to continuously capture how these tumor cells behave over hours and days. This system recreates near-physiological conditions while providing clear, dynamic visualization of invasion across different brain regions. Using this platform, we discovered that the developmental stage of brain tissue, whether it resembles that of a young child or an adolescent, has a profound impact on how aggressively pontine tumors invade. We also demonstrated that this system can effectively evaluate whether experimental drugs can block tumor invasion, providing a practical tool for screening potential therapies.

### IMPACT

This new platform addresses a critical gap in pediatric brain cancer research by enabling rapid, detailed studies of tumor invasion under conditions that closely mimic the patient’s brain environment. By revealing that the developmental stage profoundly influences tumor invasion patterns, our findings help explain why these cancers behave differently across age groups and may inform age-specific treatment strategies. Most importantly, this system provides a practical, efficient tool for screening anti-invasive therapies, a capability that has been technically challenging to achieve. Given that invasion is a primary driver of lethality in diffuse midline gliomas and current treatment options remain extremely limited, this platform could accelerate the identification of drugs that block tumor spread. By bridging the gap between basic research and therapeutic development, this approach has the potential to advance new treatment options for pediatric patients facing these devastating and currently incurable cancers.

## Data Availability Statement

All data and materials are freely available for non-commercial research upon request.

## Notes

### Competing Interest Statement

The authors have declared no competing interest.

