## Supplementary Figure and methods for "Real-time monitoring of pediatric high-grade glioma invasion using organotypic brain cultures reveals a developmentally dependent sensitivity"

### Supplementary Figure 1: Slice culture setup

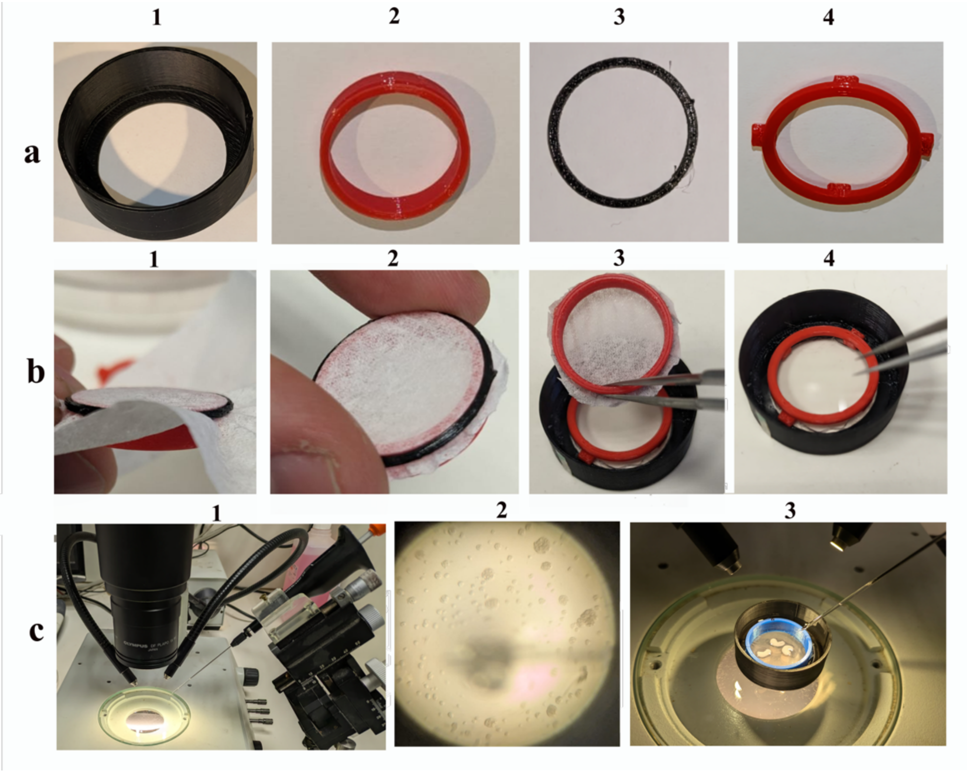

The basic elements consist of 3D-printed components, a membrane, and a glass coverslip. The coverslip is placed into a plastic holder a1 using grease to seal the chamber. To set up the stage for the slices, the membrane is positioned around a2 and secured with ring a3 (see b1). After cutting the excess membrane (b2), the membrane is positioned on stage a4 (see b3), and medium is added until it reaches the level of the membrane (b4). The membrane immediately becomes transparent upon contact with the medium, and the slices are positioned on the membrane. Twenty-four hours after plating, the injection setup is prepared (c1). Small spheroids (c2) are then picked up using a glass pipette and injected into the slices (c3). Following the injection, the medium is replaced and then refreshed every two days thereafter.

During imaging, the ring a2 is removed to minimize the distance between the objective and the slice.

**Supplementary methods:**

Patient-derived cell culture***:***

All tumor lines were maintained in "tumor stem media" (TSM), consisting of 50% neurobasal (-A) (Invitrogen), and 50% DMEM-F12 1:1 (Biological Industries) with-B27 (Without vitamin A) (Invitrogen), human bFGF (20 ng $ml^{-1}$), human EGF (20 ng $ml^{-1}$), human PDGF-AA (10 ng $ml^{-1}$) and PDGF-BB (10 ng $ml^{-1}$) and heparin (2 µg $ml^{-1}$ ; Sigma Aldrich). All growth factors were purchased from (Reprokine St Petersburg, FL, USA), as described previously [16]. All cells used were labeled with GFP or RFP as described recently [16].

All cell culture models were received from Michelle Monje (1,2) (Stanford University School of Medicine), and validated by short tandem repeat (STR) DNA fingerprinting and tested for Mycoplasma. Information about age, sex, and clinical characteristics associated with each cell culture can be found in the table below:

| Patient derived cell lines | Tumor type, location and grade | Age at diagnosis (years) | Sex | Histone – mutational status | Prior therapy |
|---|---|---|---|---|---|
| SU-DIPGXXV | DIPG, pons, WHO grade IV | 4 | F | H3.3K27M | XRT |
| SU-DIPGXIII | DIPG, pons, WHO grade IV | 6 | F | H3.3K27M | XRT |
| SU-pcGBM2 | Pediatric cortical gliobastoma, WHO grade IV | 15 | M | wt | none |

<u>Dissection and slice preparation</u>

Cortical and pontine slice cultures were prepared from 1-, 4-, or 12-week-old mice. Coronal brain slices (350 µm thick) were collected using a Leica Vibratome (VT1000s), with Campden blades (752/1/SS/50). The sections were placed on polytetrafluoroethylene 0.4 µm Pore Size (PTFE) membranes (Millipore, Sigma cat. no. BGCM00010), which become transparent when wet. The membrane was then placed on a custom setup of plastic plates and holders designed to minimize the distance between the preparation and the microscope objective. This setup incorporated a glass bottom to maximize resolution (for details, see Supplementary Figure 1). Each well was filled with medium (Neurobasal, 15 mg/ml glucose, AmphoB 25 µg/ml, Pen-

Strep, and B27 Supplement [without vitA],17504044) up to the level of the membrane to allow diffusion. Incubation was for up to 7 days in 5% CO2, 37° C. The *ex vivo* solution was replaced periodically every 1-2 days.

Immunofluorescence:

Deparaffinization was performed using xylene washes, followed by a gradient of ethanol washes (100%-50%). Antigen retrieval was conducted using a citrate buffer (10 mM sodium citrate in 0.055% Tween-20) at 95°C for 15 minutes. Sections were permeabilized with a buffer (0.2% Triton X-100 in PBS) for 45 minutes, followed by blocking for 1 hour in 10% BSA in PBS. Primary antibodies were applied overnight at 4°C, followed by secondary antibodies for 1 hour. Sections were then mounted using Fluoroshield mounting medium containing DAPI (Abcam,cat no.104139) .

List of Antibodies:

Chicken anti - GFP (Abcam, Cat No. ab13970)

Rabbit anti - CD11b (CST, 93169) Cat No.17198

Rabbit anti - ALDH1L1 (Abcam, Cat No. ab87117)

Rabbit anti - NeuN (Cat No. 12943)

Rabbit anti - Ki67 (Thermo, RM-9106-s1)

Slice preparation and staining

Slices were fixed in 4% formalin (Bio-Lab, Israel Cat. No. 0005450305F1) overnight at 4°C, then transferred to 2% UltraPure™ Low Melting Point Agarose (Cat. No. 16520-050 Thermofisher Scientific, Waltham, MA, USA). The embedded tissue was then transferred to HISTOSETTE® I Tissue Processing/Embedding Cassettes (cat. No. H0542, Merck, Sigma). The cassettes were transferred to 50% ethanol for 1 hour, then to 80% ethanol, where they were kept until further processing. Cassettes were processed in a Magnus device (Milestone) according to a standard protocol. Blocking and embedding in paraffin were performed using a HistoCore Arcadia-C device (Leica), followed by sectioning on a microtome (Leica RM2255) and drying in a 60°C oven overnight. H&E staining was performed using a cover slipper device (Sakura Tissue-Tek film).
